# The Quantification of Antibody Elements and Receptors subunit expression using qPCR: The Design of VH, VL, CH, CL, FcR subunits primers for a more holistic view of the immune system

**DOI:** 10.1101/714550

**Authors:** Wei-Li Ling, Yuen-Ling Ng, Anil Wipat, David Philip Lane, Samuel-Ken-En Gan

## Abstract

The expression levels of Immunoglobulin elements and their receptors are important markers for health and disease. Within the immunoglobulin locus, the constant regions and the variable region families are associated with certain pathologies, yet a holistic view of the interaction between the expression of the multiple genes remain to be fully characterized. There is thus an important need to quantify antibody elements, their receptors and the receptor subunits in blood (PBMC cDNA) for both screening and detailed studies of such associations. Leveraging on qPCR, we designed primers for all Vκ 1-6, VH1-7, Vλ1-11, nine CH isotypes, Cκ, Cκ, Cλ1 &3, FcεRI α,β, and γ subunits, all three FcγR and their subunits, and FcαR. Validating this on a volunteer PBMC cDNA, we show a qPCR primer set repertoire that can quantify the relative expression of all the above genes to GAPDH housekeeping gene, with implications and uses in both clinical monitoring and research.

## 1. Introduction

There are large numbers of immune genes of the adaptive immune response and their expressions levels have been associated to various states of health and disease. In healthy individuals, the ratio of expression of these immune genes (e.g. antibodies) can be associated with pre-disease states. Biases in Immunoglobulin (Ig) variable heavy chains (VH chains) 1, 3, and 4 families, were found to be more prominent in asthma (Snow et al., 1995), with VH1 reported to be dominant in patients suffering from peanut allergy (Janezic et al.,1998). High levels of certain antibody isotypes were also found to be associated with specific diseases e.g. IgE for allergy (Stanworth., 1993), IgA in IgA nephropathy (Suzuki, H., 2011), and IgM in hyper-IgM syndrome (Lougaris, V., 2005) amongst many others. The constant region (forming the isotype) was also able to influence the ability of V-regions to bind antigens (Lua et al., 2018) and the variable light chain (VL) families were also found to impact the interaction with antigens (Ling et al., 2017), production (Ling et al., 2017; Ling et al., 2018), and the interaction with Ig receptors (FcRs; see Ling et al., 2018; Su et al., 2018; Lua et al., 2019). Given the effects of the various antibody parts and their association with certain pathologies, the relative quantification of specific antibody elements and genes together with the antibody receptors and their subunits are important markers in health screening and to provide a more holistic investigation to disease associations. Since antibodies often work in tandem with effector immune cells via the FcR, quantifying FcR expression levels in PMBC provide an insight to the responses available to the various antibody isotypes. To address this, we aim to develop a set of primers for qPCR to study the spectrum of immune genes in PBMCs. The method developed here aims to aid the diagnostic and further study of immunological conditions and disease.

Our primer repertoire targets the six kappa variable light chains families (Vκ1-6). eleven lambda variable light chains families (Vλ1-11), seven variable heavy chain families (VH1-7), one kappa constant light chain (Cκ), seven lambda constant light chains (Cλ 1-7), and nine constant heavy chains (Ig γ 1-4, Igα1&2, Igε, Igμ and Igδ), along with the respective Fc receptor subunits: FcγR1α, 2α, 2β, 2c, 3α, 3β, FcαR, FcεR1α, FcεR1β, FcεR1γ & FcεR2. We quantified the target messenger ribonucleic acid (mRNA) through complementary deoxyribonucleic acid (cDNA) synthesis using quantitative real-time polymerase chain reaction (qPCR) and showed their levels in human PBMC cDNA.

## 2. Material and methods

### 2.1 Designing qPCR primers

Full receptor DNA sequences were obtained from GenBank^®^ (https://www.ncbi.nlm.nih.gov/genbank, Benson et al., 2013), and immunoglobulin DNA sequences were obtained from IMGT (http://www.imgt.org, Lefranc et al., 1999), with the following accession numbers shown in Table A.1. DNA sequences of target genes were analysed using the Primer3plus program (http://www.bioinformatics.nl/cgibin/primer3plus/primer3plus.cgi, Untergasser et al., 2007) for qPCR. Primer output were subjected to UCSC In-Silico PCR program (https://genome.ucsc.edu/cgi-bin/hgPcr, Kuhn et al., 2013) for verification of specificity on the target gene.

### 2.2. Real-Time Polymerase Chain Reaction (qPCR)

The quantification of gene expression was obtained using Applied Biosystem StepOnePlus™. PowerUp™ SYBR® Green Master Mix (2X) reagents according to the manufacturer’s recommendations. Each gene target quantification reaction was performed separately with the respective primer sets. For simultaneous operation, the amplification and melt curve reaction settings utilized a constant annealing temperature despite the different theoretical melting temperatures of each primer set. The settings are as follows: Stage 1 (1 cycle) at 50°C, for 120 seconds, followed by 95°C for 2 seconds; Stage 2 (40 cycles) at 95°C for 3 seconds, followed by 60°C for 30 seconds; and Melt curve at 95°C for 15 seconds followed by 60°C for 60 seconds, and finally 95°C for 15 seconds. Optimization and validation of primer pairs consisted of triplicate independent runs with technical triplicates including “No Template Controls” (NTC) and samples for each run.

### 2.3. Gene Expression Levels

Gene expression results were analysed using Applied Biosystem software, StepOnePlus™ Version 2.3. Threshold Cycle (CT) values were auto-populated with all parameters set on default with Glyceraldehyde 3-phosphate dehydrogenase (GAPDH) housekeeping gene as the endogenous control. The GAPDH gene was used as baseline to deduct each gene expression CT mean value from triplicate independent runs (including technical triplicates), further divided by GAPDH gene CT mean value and converted to percentage using the following formula: 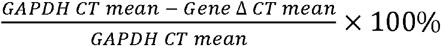

## 3. Results and discussion

### 3.1. Primer design

Since antibodies are formed by V(D)J recombination rather than by the artificial VH and VL frameworks (discussed in Phua et al., 2019), many antibody target genes have several different germline sequences classified under same VH or VL family. It is thus difficult to design primers for each individual germline sequence. To overcome this issue, we selected the representatives of each target gene (of the VH and VL families) through sequence alignment of all other germlines to determine the most conserved regions. The resulting list of qPCR primers was generated based on Vκ families (Table 1), VH families (Table 2), Vλ families (Table 3), CH, and both CL: Cκ and Cλ (Table 4) alongside the immunoglobulin receptor subunits and GAPDH (Table 5) using the Primer3plus program (refer to 2.1)

**Table 1.**
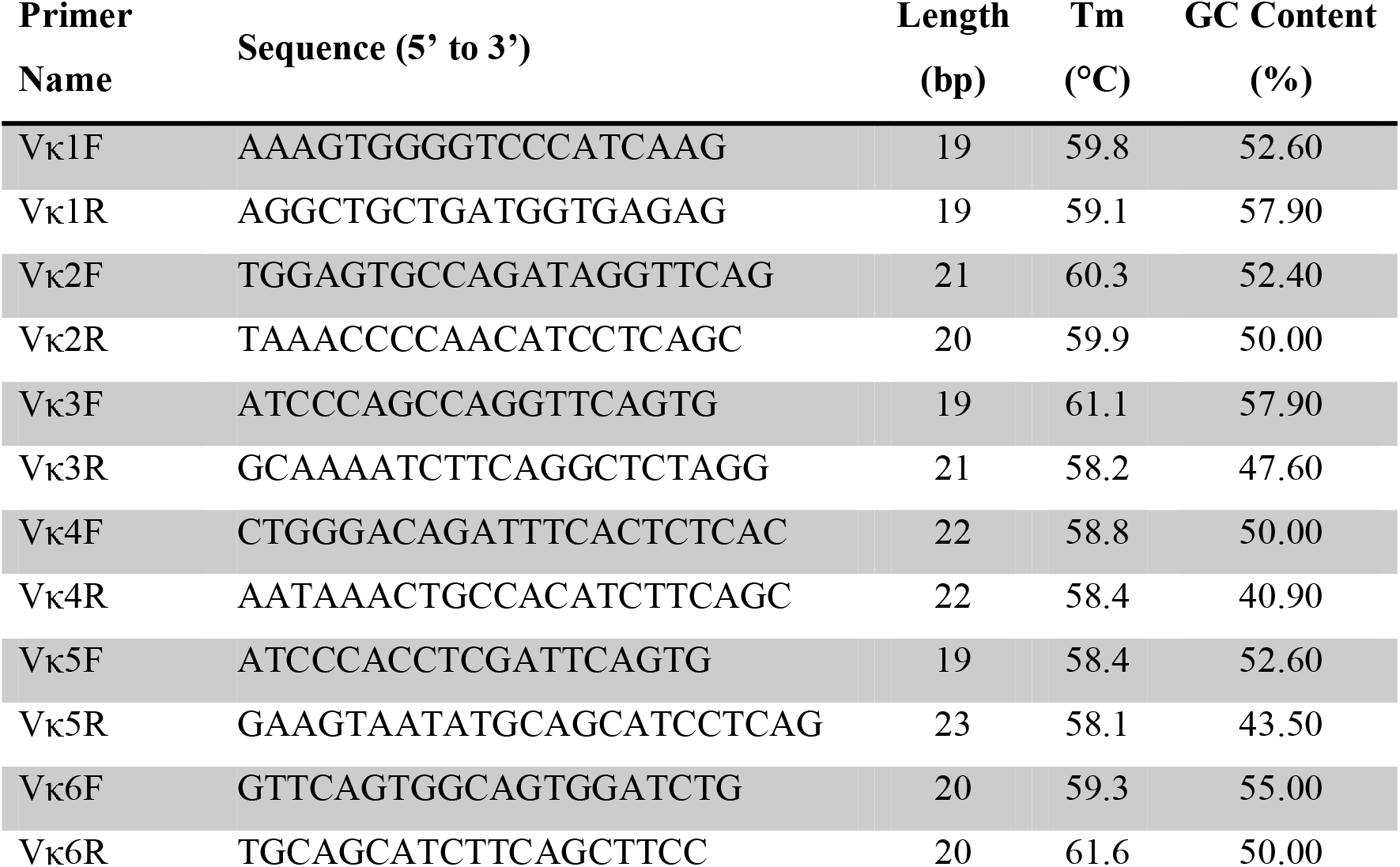
List of qPCR forward and reverse primers presented in 5’ to 3’ fashion generated using Primer3plus program against Vκ families as gene targets with respective primer lengths, melting temperatures and GC contents listed.

**Table 2.**
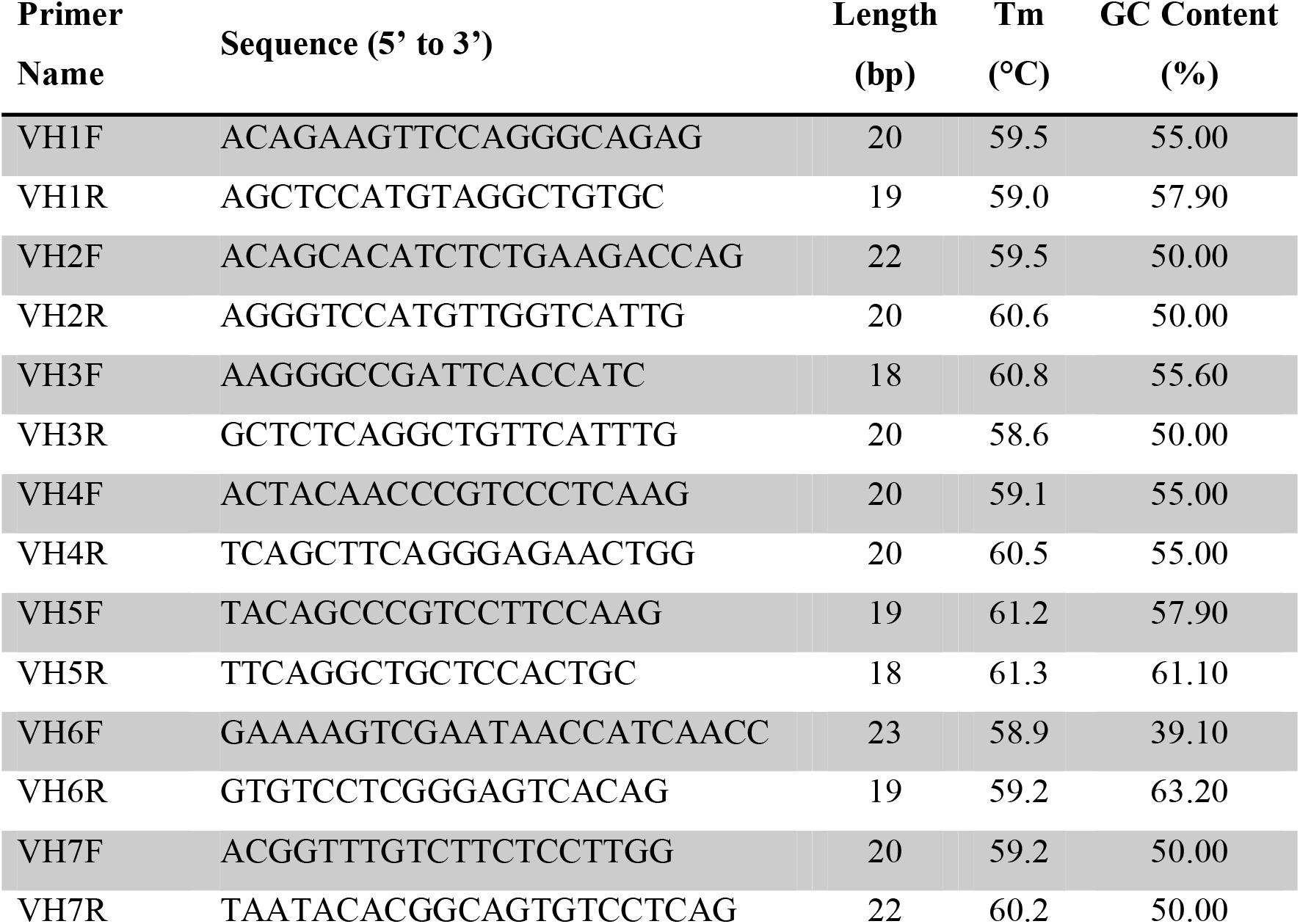
List of qPCR forward and reverse primers presented in 5’ to 3’ fashion generated using Primer3plus program against VH families as gene targets with respective primer lengths, melting temperatures and GC contents listed.

**Table 3.**
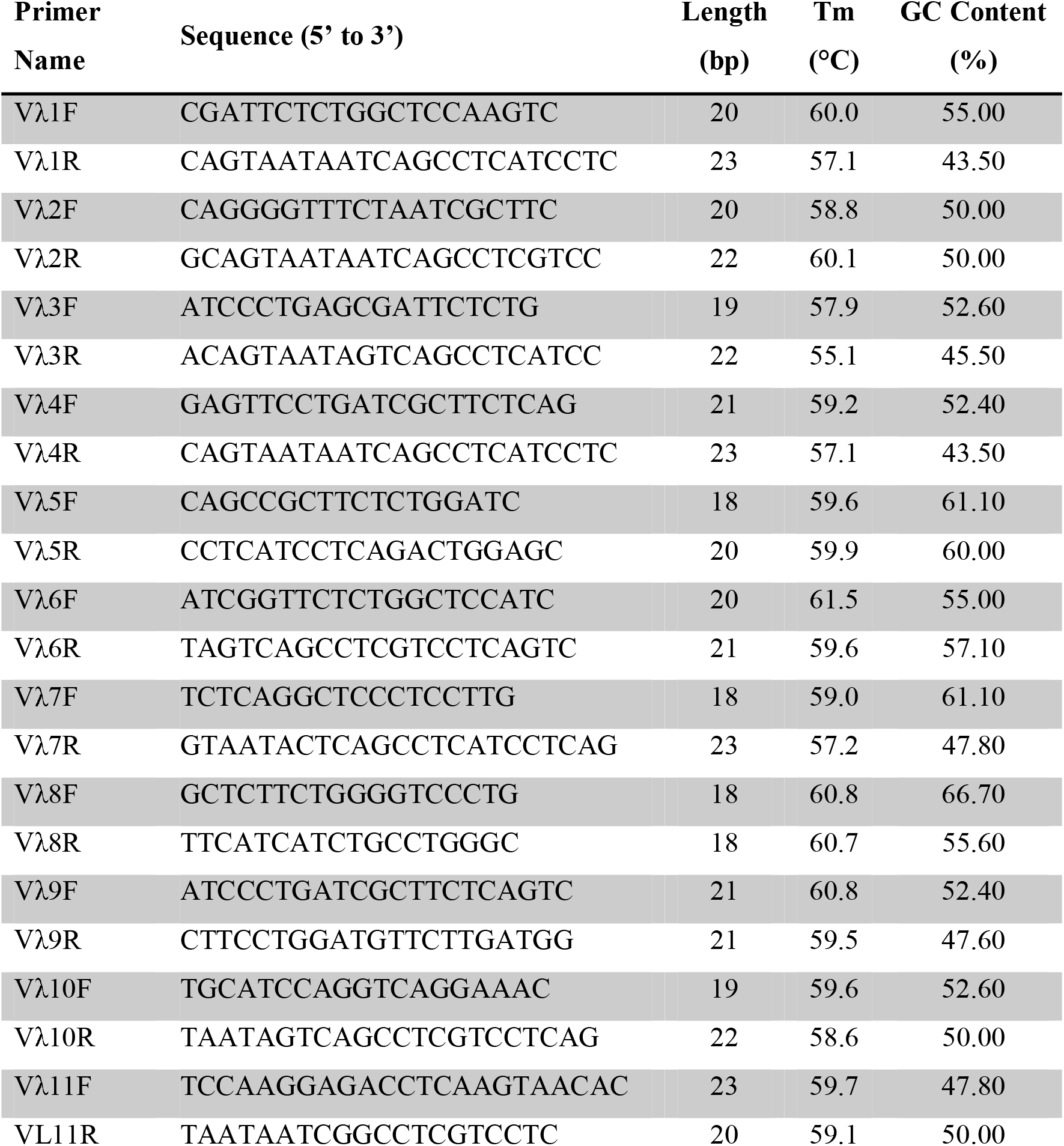
List of qPCR forward and reverse primers presented in 5’ to 3’ fashion generated using Primer3plus program against Vλ, families as gene targets with respective primer lengths, melting temperatures and GC contents listed.

**Table 4.**
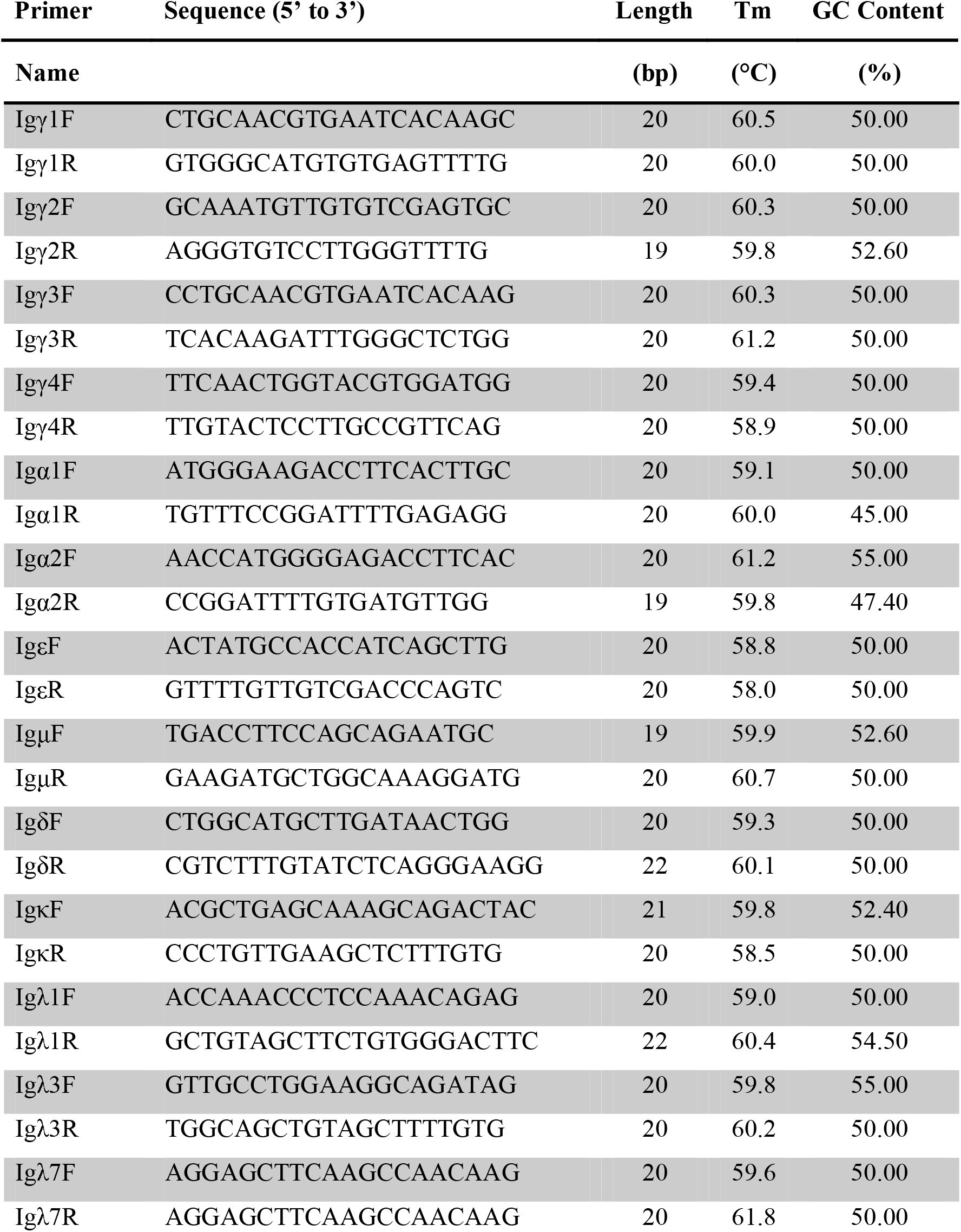
List of qPCR forward and reverse primers presented in 5’ to 3’ fashion generated using Primer3plus program against CHs, Cκ and Cλs as gene targets with respective primer lengths, melting temperatures and GC contents listed.

**Table 5.**
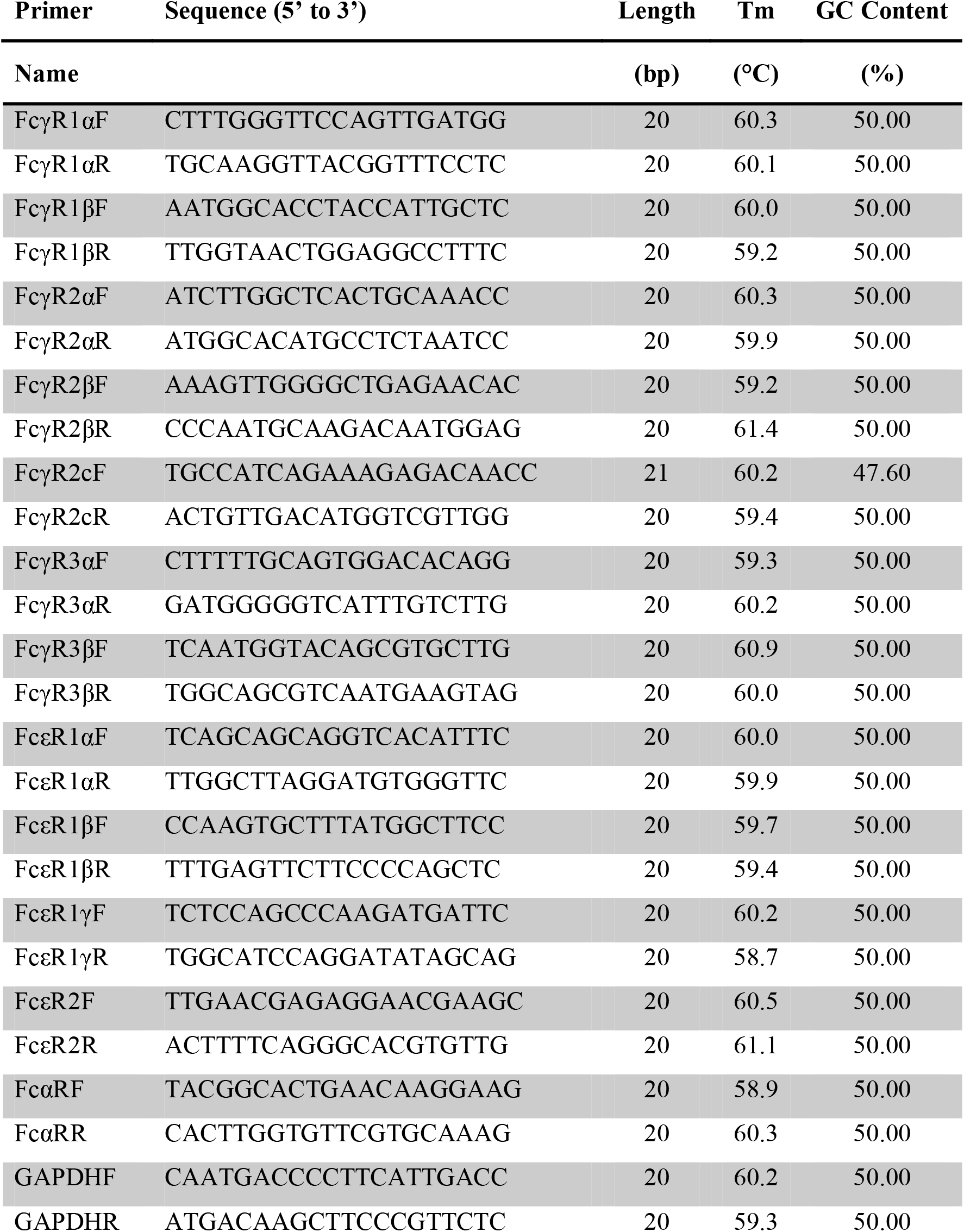
List of qPCR forward and reverse primers presented in 5’ to 3’ fashion generated using Primer3plus program against immunoglobulin receptor subunits and GAPDH as gene targets with respective primer lengths, melting temperatures and GC contents listed.

For the primers to amplify the correct gene targets in the pool of cDNA, the specificity is important. However, there are a few highly related gene targets such as the Cλ 2, 4, 5, 6 and 7 that complicated the design of specific primers against these targets, and thus they were excluded.

Primers were chosen with melting temperature (Tm) as close to 60°C and GC content around 50% with amplicon product sizes between 70 to 150 base pair (bp).

Since FcαμR (with dual specificity to antibody isotype IgA and IgM), is only expressed and found in niche populations of IgD+/CD38+ mature B cells in tonsil tissue (Kikuno et al., 2007) and activated macrophages (Feng et al., 2010), it was thus excluded from the design (Table 5) given that the purpose was focused towards blood PBMC cDNA quantification.

### 3.2. Threshold cycle (Ct) data of gene targets

After designing the primers, we validated them on PBMCs. Three independent runs with three technical replicates, Ct mean values, delta Ct mean and delta Ct standard error (SE) were calculated as shown in Table 6.

**Table 6.** CT value of various gene targets populated from StepOnePlus™ Version 2.3. Gene target GAPDH were used as endogenous control as well as normalization. CT mean, delta CT mean and CT SE values are obtained from independent triplicate run with each run containing triplicate technical samples.

| Gene Target | Acronym | Ct Mean | ΔCt Mean | ΔCt SE |
| --- | --- | --- | --- | --- |
| Glyceraldehyde 3-phosphate dehydrogenase | GAPDH | 16.23 | - | - |
| Variable Heavy 1 | VH1 | 23.14 | 6.58 | 0.05 |
| Variable Heavy 2 | VH2 | 26.04 | 9.48 | 0.07 |
| Variable Heavy 3 | VH3 | 17.17 | 0.61 | 0.07 |
| Variable Heavy 4 | VH4 | 21.78 | 5.22 | 0.05 |
| Variable Heavy 5 | VH5 | 21.78 | 5.22 | 0.05 |
| Variable Heavy 6 | VH6 | 28.32 | 11.76 | 0.05 |
| Variable Heavy 7 | VH7 | 27.09 | 10.53 | 0.06 |
| Variable Light Kappa 1 | Vκ1 | 17.78 | 1.23 | 0.06 |
| Variable Light Kappa 2 | Vκ2 | 22.37 | 5.81 | 0.05 |
| Variable Light Kappa 3 | Vκ3 | 24.24 | 7.68 | 0.06 |
| Variable Light Kappa 4 | Vκ4 | 25.17 | 8.62 | 0.05 |
| Variable Light Kappa 5 | Vκ5 | 30.79 | 14.24 | 0.08 |
| Variable Light Kappa 6 | Vκ6 | 28.53 | 11.98 | 0.05 |
| Variable Light Lambda 1 | Vλ1 | 19.54 | 2.98 | 0.05 |
| Variable Light Lambda 2 | Vλ2 | 24.27 | 7.69 | 0.1 |
| Variable Light Lambda 3 | Vλ3 | 20.10 | 3.66 | 0.41 |
| Variable Light Lambda 4 | Vλ4 | 22.58 | 5.99 | 0.12 |
| Variable Light Lambda 5 | Vλ5 | 28.04 | 11.45 | 0.07 |
| Variable Light Lambda 6 | Vλ6 | 27.4 | 10.81 | 0.05 |
| Variable Light Lambda 7 | Vλ7 | 26.42 | 9.83 | 0.04 |
| Variable Light Lambda 8 | Vλ8 | 25.7 | 9.12 | 0.04 |
| Variable Light Lambda 9 | Vλ9 | 30.33 | 13.75 | 0.04 |
| Variable Light Lambda 10 | Vλ10 | 27.21 | 10.63 | 0.07 |
| Variable Light Lambda 11 | Vλ11 | 35.79 | 19.16 | 0.31 |
| Constant Heavy Gamma 1 | Igγ1 | 18.55 | 1.97 | 0.04 |
| Constant Heavy Gamma 2 | Igγ2 | 23.06 | 6.48 | 0.05 |
| Constant Heavy Gamma 3 | Igγ3 | 28.4 | 11.81 | 0.04 |
| Constant Heavy Gamma 4 | Igγ4 | 16.97 | 0.39 | 0.12 |
| Constant Heavy Alpha 1 | Igα1 | 20.83 | 4.81 | 0.05 |
| Constant Heavy Alpha 2 | Igα2 | 21.73 | 5.71 | 0.04 |
| Constant Heavy Delta | Igδ | 23.89 | 7.87 | 0.06 |
| Constant Heavy Epsilon | Igε | 27.31 | 11.29 | 0.06 |
| Constant Heavy Mu | Igμ | 19.87 | 3.86 | 0.06 |
| Constant Light Kappa | IgK | 19.02 | 3.09 | 0.04 |
| Constant Light Lambda 1 | Igλ1 | 20.14 | 4.22 | 0.04 |
| Constant Light Lambda 3 | Igλ3 | 17.74 | 1.82 | 0.11 |
| Fc Alpha Receptor | FcαR | 26.4 | 10.47 | 0.03 |
| Fc Epsilon Receptor 1 Alpha | FcεRIα | 23.87 | 7.85 | 0.06 |
| Fc Epsilon Receptor 1 Beta | FcεRIβ | 27.92 | 11.91 | 0.06 |
| Fc Epsilon Receptor 1 Gamma | FcεRIγ | 21.42 | 5.49 | 0.06 |
| Fc Epsilon Receptor 2 | FcεRII | 25.43 | 9.5 | 0.03 |
| Fc Gamma Receptor 1 Alpha | FcγRIα | 27.56 | 11.55 | 0.07 |
| Fc Gamma Receptor 1 Beta | FcγRIβ | 28.61 | 12.6 | 0.05 |
| Fc Gamma Receptor 2 Alpha | FcγRIIα | 24.2 | 8.19 | 0.1 |
| Fc Gamma Receptor 2 Beta | FcγRIIβ | 26.64 | 10.63 | 0.04 |
| Fc Gamma Receptor 2 c | FcγRIIc | 26.2 | 10.18 | 0.06 |
| Fc Gamma Receptor 3 Alpha | FcγRIIIα | 25.8 | 9.78 | 0.06 |
| Fc Gamma Receptor 3 Beta | FcγRIIIβ | 29.33 | 13.31 | 0.04 |

The primer sets were evaluated based on the returned Ct mean values generated from the StepOnePlus™ software. As shown in Table 6, all primer sets had acceptable returned values ranging from the lowest Ct value of 16.23 (GAPDH) to the highest Ct value of 30.79 (Vĸ5). One exception was the Vλ1 (Ct value of 35.79 that is near to the end of total amplification cycles) gene that had either insufficient template or unspecific amplification. The latter was more likely since the values were populated from three independent runs with technical triplicates with low Ct standard error (SE), suggesting that the expression levels of Vλ11 to be of sufficient amounts for the assay. It should be noted that despite having a return CT mean value, Vλ11 expression levels in the volunteer were measured to be close to 0% (Table 6).

### 3.3. Gene expression level of gene targets

In order to compare between the gene targets, the relative comparison method was deployed using GAPDH as a standard (relative percentage, 100%) among all independent runs and technical triplicates. Our results (Figure 1) showed that the genes had expected levels of expression. VH3 and Vκ1 were expectedly more abundant amongst the VH and Vκ families, a finding in agreement with a previous report (Tiller et al., 2013) There was also expected variations within the CLs, although the Cκ and Cλ levels were relatively balanced in our data and not in agreement with previous findings (Haraldsson et al., 1991; Normansell., 1987). Nonetheless, this can be explained by volunteer factors. Another unexpected ratio is the IgG4, IgE and IgD levels of the volunteer. While IgG4 should be at around 4% the total amount of IgG (Hashira et al., 2000; Janeway et al., 2001), the volunteer IgG4 levels exceeded the expression levels of IgG1. Similarly, the volunteer had a 30% IgE expression level when the typical IgE level in healthy individuals should be extremely low at ~ 0.000175% of the total immunoglobulin levels, and certainly lower than that of IgG4. Since the volunteer has a history of severe Type-I hypersensitivity, this result may not be unexpected, and in fact, supported previous reports that the ratio between IgG4 and IgE are markers for allergy (Caubet et al., 2012; Noh et al., 2007). For IgD, the volunteer had an elevated level at 52% while it is typically about 0.23% of the total serum immunoglobulin (Janeway et al., 2001). Regardless, much remains elusive when it comes to IgD, although some studies show that IgD can replace IgM in IgM immunodeficiency (Lutz et al., 1998), which may not explain our findings as the individual did not suffer from IgM immunodeficiency given the volunteer’s IgM levels of 76%. Such discrepancies demonstrate the need to study the IgM/IgD ratios and that of the various genes, especially at various health and disease states for a more holistic understanding.

**Figure 1.**
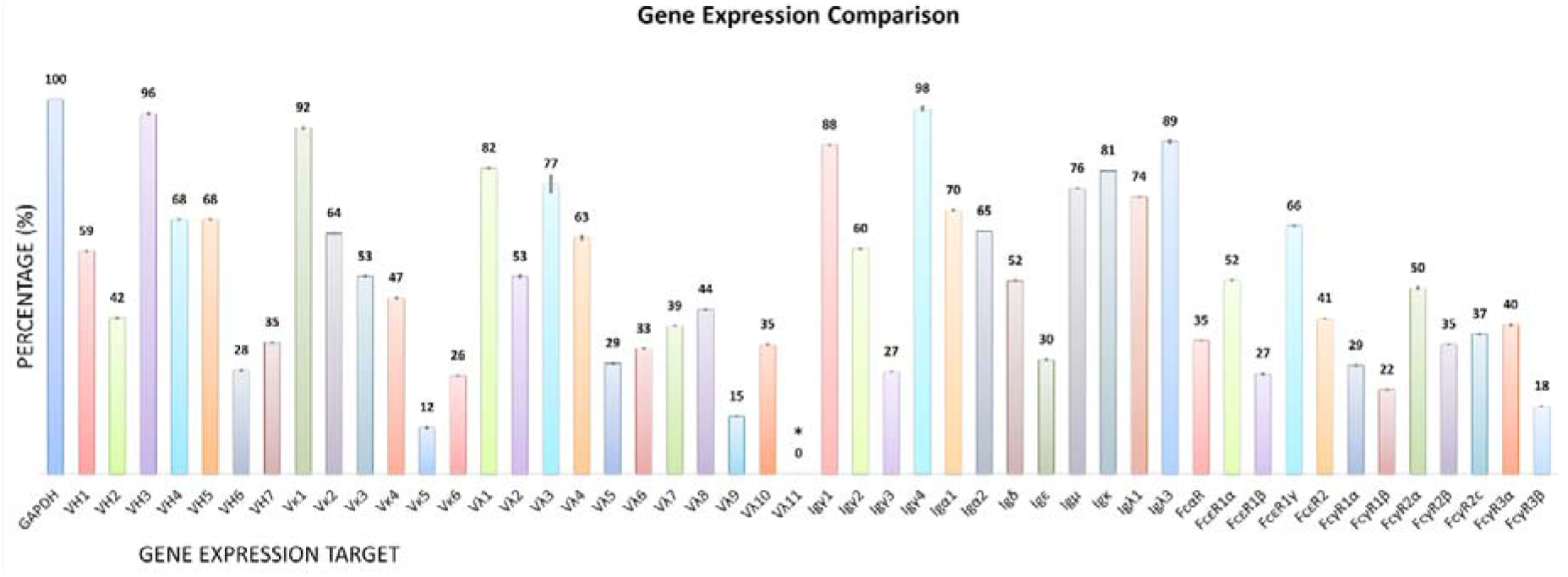
The Histogram showing various gene expression percentages. The housekeeping GAPDH was used as an endogenous control for normalization to 100%. Percentage were calculated by subtracting delta CT mean value of each gene from GAPDH CT mean value followed by division of GAPDH CT mean value and multiplied by 100% (refer to 2.1). Asterisk (*) is a percentage value being replaced instead of original (−18%) due to ACT mean reading.

## 4. Conclusion

We set out to design primers that could detect the various V-region families of antibodies, the antibody isotypes and subtypes, as well receptors subunits to quantify their relative expression in PBMCs. Leveraging on the use of qPCR, the quantification could be performed with very little amounts of blood such as those extracted from finger pricks (Poh et al., 2014). By combining finger prick blood methods and qPCR primers, screening and diagnosis in the clinical setting can be performed easily. In the research front, a more holistic view of the expression ratios can be gathered to shed light on their association with health and disease.

## Author Contributions

WWL performed the experiments, analysis and drafted the manuscript. YLN, AW, DPL were involved in the writing and discussions of the manuscript. SKEG conceived and supervised all aspects of the study.

## Appendices

**Table A.1.** List of genes with corresponding identify or reference number used in the designing of qPCR primers.

| **Gene Name** | **Identification Reference / Number** |
| --- | --- |
| VK1 | IMGT Reference Sequence: <u>**V01577**</u> |
| VK2 | IMGT Reference Sequence: <u>**M31952**</u> |
| VK3 | IMGT Reference Sequence: <u>**X12686**</u> |
| VK4 | IMGT Reference Sequence: <u>**Z00023**</u> |
| VK5 | IMGT Reference Sequence: <u>**X02485**</u> |
| VK6 | IMGT Reference Sequence: <u>**X63399**</u> |
| VH1 | IMGT Reference Sequence: <u>**Z14296**</u> |
| VH2 | IMGT Reference Sequence: <u>**L21969**</u> |
| VH3 | IMGT Reference Sequence: <u>**M99660**</u> |
| VH4 | IMGT Reference Sequence: <u>**M99684**</u> |
| VH5 | IMGT Reference Sequence: <u>**996686**</u> |
| VH6 | IMGT Reference Sequence: <u>**J04097**</u> |
| VH7 | IMGT Reference Sequence: <u>**L10057**</u> |
| VL1 | IMGT Reference Sequence: <u>**M94116**</u> |
| VL2 | IMGT Reference Sequence: <u>**Z73664**</u> |
| VL3 | IMGT Reference Sequence: <u>**X97473**</u> |
| VL4 | IMGT Reference Sequence: <u>**Z73648**</u> |
| VL5 | IMGT Reference Sequence: <u>**Z73670**</u> |
| VL6 | IMGT Reference Sequence: <u>**Z73673**</u> |
| VL7 | IMGT Reference Sequence: <u>**Z73674**</u> |
| VL8 | IMGT Reference Sequence: <u>**Z73650**</u> |
| VL9 | IMGT Reference Sequence: <u>**Z73675**</u> |
| VL10 | IMGT Reference Sequence: <u>**Z73676**</u> |
| VL11 | IMGT Reference Sequence: <u>**D86996**</u> |
| IgG1 | IMGT Reference Sequence: <u>**J00228**</u> |
| IgG2 | IMGT Reference Sequence: <u>**J00230**</u> |
| IgG3 | IMGT Reference Sequence: <u>**D78345**</u> |
| IgG4 | IMGT Reference Sequence: <u>**K01316**</u> |
| IgA1 | IMGT Reference Sequence: <u>**J00220**</u> |
| IgA2 | IMGT Reference Sequence: <u>**J00221**</u> |
| IgE | IMGT Reference Sequence: <u>**J00222**</u> |
| IgM | IMGT Reference Sequence: <u>**X14940**</u> |
| IgD | IMGT Reference Sequence: <u>**X57331**</u> |
| FcgR1a | NCBI Reference Sequence: <u>**NM_000566**</u> |
| FcgR1b | NCBI Reference Sequence: <u>**NM_001244910**</u> |
| FcgR2a | NCBI Reference Sequence: <u>**NM_021642**</u> |
| FcgR2b | NCBI Reference Sequence: <u>**NM_001190828**</u> |
| FcgR2c | NCBI Reference Sequence: <u>**NR_047648**</u> |
| FcgR3a | NCBI Reference Sequence: <u>**NM_000569**</u> |
| FcgR3b | NCBI Reference Sequence: **<u>NM_000570</u>** |
| FceR1a | NCBI Reference Sequence: **<u>NM_002001.3</u>** |
| FceR1b | NCBI Reference Sequence: **<u>NM_000139</u>** |
| FceR1g | NCBI Reference Sequence: **<u>NM_004106.2</u>** |
| FceR2 | NCBI Reference Sequence: **<u>NM_001207019</u>** |
| FcaR | NCBI Reference Sequence: **<u>NM_133269</u>** |
| GAPDH | NCBI Reference Sequence: **<u>NM_002046</u>** |

